# Accumbens D2-MSN hyperactivity drives behavioral supersensitivity

**DOI:** 10.1101/2020.11.12.380667

**Authors:** Anna Kruyer, Jeffrey Parilla-Carerro, Courtney Powell, Lasse Brandt, Stefan Gutwinski, Ariana Angelis, Reda Chalhoub, Davide Amato

## Abstract

Antipsychotic-induced behavioral supersensitivity is a problematic consequence of long-term treatment with antipsychotic drugs and is characterized by emergence of refractory symptoms and dyskinesias. The underlying mechanisms are unknown, and no rational approaches exist to prevent or reverse antipsychotic-induced supersensitivity. Here we describe major adaptations impacting populations of striatal medium spiny neurons (MSNs) during the development of behavioral supersensitivity and reveal a prominent role played by D2 receptor expressing MSNs. We show that enhanced D2-MSN activity underlies several symptoms spanning from psychostimulant sensitization, to antipsychotic treatment resistance and drug addiction. Our data warn against severe adverse events following antipsychotic treatment discontinuation and offer insight that may inform therapeutic approaches to overcome antipsychotic-induced supersensitivity.

## Introduction

Antipsychotic drugs are widely prescribed for psychotic and non-psychotic disorders (*1–3*). Patients with psychotic symptoms often discontinue antipsychotic treatment due to diminished therapeutic efficacy and emergence of problematic side effects (*4*). Antipsychotic discontinuation may exacerbate refractory symptoms (*5–7*) and pose a risk for developing tardive dyskinesia (*6, 8–10*) and substance use disorder (*11, 12*). These potential outcomes, referred to as behavioral supersensitivity are hypothesized to result from enhanced sensitivity to dopamine via upregulated D2 receptors (D2r) owing to long-term receptor blockade (*13*), even though increases in D2r expression, binding affinity, and/or function are not consistently reported in humans or animal models during or after treatment with clinically relevant doses of antipsychotics (*5, 14*). In previous studies we described pre- and post-synaptic changes in neurotransmission during antipsychotic treatment resistance despite adequate D2r blockade (*15*). We also found that discontinuation from chronic antipsychotic treatment alters the physiology of striatal cells (*16*), but we did not define specific adaptation patterns nor their significance for behavioral symptoms. Here we describe characteristic symptoms and the underlying neurobiology that stem from discontinuing chronic antipsychotic after treatment failure (*15*) in both mice and rats.

### Antipsychotic discontinuation and motor side effects in animal models

Whether treatment discontinuation following chronic antipsychotic regimens causes motor side effects such as dyskinesia is unclear since studies report conflicting results using animal models (*17–20*). We performed a systematic review of the literature on spontaneous vacuous chewing movements (VCM) in rodents, which are considered a proxy for extrapyramidal and/or oral dyskinesia symptoms induced by antipsychotic drug treatment (*17*). We searched PubMed, EMBASE, and Web of Science for relevant studies using search terms for (1) animal studies, (2) antipsychotics, and (3) withdrawal (See Supplemental material). We found that discontinuation from second generation antipsychotic drugs induces no spontaneous VCMs, with the exception of high doses of risperidone, whereas haloperidol induces high levels of spontaneous VCMs in young and old rats 21 days after prolonged daily treatment (75d) (fig. S1). To determine if abstinence from shorter treatment regimens using haloperidol caused oral dyskinesia, we measured VCMs in rats 7d after haloperidol discontinuation, but did not observe VCMs using this paradigm (fig. S2) in keeping with reports using similar treatment durations (*21*).

Brief abstinence (72h) from high doses of daily repeated i.p. haloperidol (15d) results in hyperlocomotion induced by cocaine (*22*), consistent with behavioral supersensitivity following antipsychotic treatment discontinuation (9). In addition, we previously found that 14d of continuous treatment with clinically-relevant doses of haloperidol results in loss of therapeutic efficacy relative to shorter treatment durations (15). To confirm whether abstinence from antipsychotics following treatment failure produced behavioral supersensitivity, we treated mice with a systemic injection of cocaine 7 days after discontinuation of 14d haloperidol treatment, delivered continuously through osmotic pumps (Fig. 1A). Acute cocaine injection in these animals enhanced locomotion (cross-sensitization) in a similar manner as in haloperidol-naïve animals receiving a second cocaine injection 7 days after an initial cocaine injection (mono-sensitization, Fig. 1B) (*23*), thus demonstrating behavioral supersensitivity.

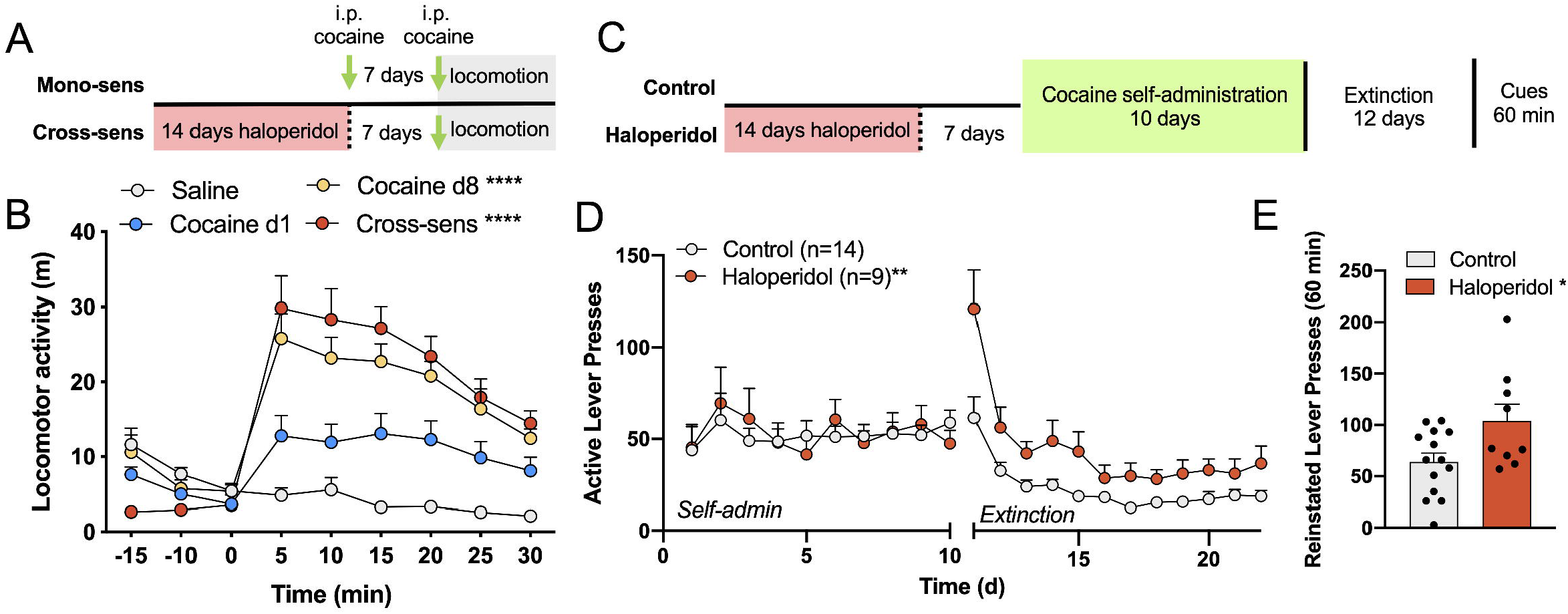

### Haloperidol discontinuation in the absence of schizophrenia produced endophenotypes of substance use disorder

The likelihood of substance use disorder in patients with schizophrenia is ~4.6-fold higher than in the general population (*24, 25*), especially among treatment non-adherent patients (*26*). Interestingly, non-psychotic drug-addicted patients are reported to co-abuse antipsychotic medications to enhance the effects of addictive substances (*27*). This raises the question of whether antipsychotic treatment itself, in the absence of psychiatric illness, enhances vulnerability to substance use disorder. Traditional animal models of drug addiction vulnerability involve locomotor sensitization induced by intermittent psychostimulant injections (*28–30*). Because haloperidol and cocaine produced locomotor cross-sensitization, we sought to determine whether this response was predictive of additional behavioral features of substance use disorder using the self-administration, extinction, and reinstatement model of cocaine use and relapse.

It has been shown previously that intake of substances of abuse is suppressed during ongoing antipsychotic treatment (*31, 32*), although lower antipsychotic doses can produce opposite outcomes (*33*). Regardless, it is unknown if discontinuing antipsychotic medications after loss of therapeutic efficacy is a risk factor for substance use disorder. To examine this hypothesis, we trained male and female rats to self-administer cocaine after abstinence from haloperidol according to the same timeline applied during locomotor cross-sensitization (Fig. 1C). Operant training was conducted on an FR1 schedule, and cocaine delivery was paired with light and tone cues for 2h each day. There were no differences in active or inactive lever pressing between haloperidol pretreated rats and control animals during self-administration and groups did not differ in total cocaine intake (Fig. 1D, fig. S3A), suggesting that the acquisition of learned operant responding and reward associated with cocaine delivery were not altered by haloperidol discontinuation. After 10 days of operant training, rats were then extinguished to the drug-associated context during 2h sessions in the operant box without cues or drug delivery. In control animals, active lever presses gradually diminished over time in the absence of drug reward and a stable baseline of responding was observed within 3-5 days of extinction training. In haloperidol-pretreated rats however, active lever pressing was elevated compared to controls throughout extinction training and baseline responding in the drug-paired context remained elevated despite absence of reward in haloperidol pretreated rats (Fig. 1D). After 12 days of extinction, animals were returned to the operant box and light/tone pairings were restored to the active lever, but no cocaine was delivered. Cocaine-associated cues stimulated lever pressing in haloperidol-pretreated rats compared to control animals during a 60-min reinstatement test (Fig. 1E), a widely accepted model of cue reactivity (*34*) linked to drug relapse in human patients (*35*). Thus the main deficit emerging in supersensitive animals was surprisingly not an increase in cocaine intake as reported in studies applying mono-sensitization procedures (*28*), but involved disrupted extinction of operant responding (i.e. lever pressing) for cocaine despite its absence. Similarly, haloperidol pretreated animals displayed enhanced measures of seeking in response to drug-associated cues, perhaps owing to a deficit in within-session extinction of behavioral responding in the absence of cocaine. Together, these data indicate that antipsychotic discontinuation produced features of substance use disorder coincident with behavioral supersensitivity.

### D2-MSN hyperactivity, but not D2 receptor upregulation, drives behavioral supersensitivity

The most commonly cited cause of antipsychotic-induced behavioral supersensitivity is upregulated D2r expression after long-term treatment (*13*). To test whether the D2r was upregulated in our model of behavioral supersensitivity, we examined D2r levels in the ventral striatum, dorsal striatum, and midbrain by Western blot after 14d of continuous haloperidol treatment and after 7d of abstinence from continuous treatment. We found no change in D2r expression in either group using our treatment protocols relative to untreated animals (fig. S4).

Spontaneous and sensitized locomotion and goal-directed behaviors are orchestrated by activity of D1 r and D2r-expressing MSNs in the ventral striatum (D1- and D2-MSNs) (*36*). To examine the cellular basis of locomotion during behavioral supersensitivity, we recorded single cell Ca^2+^ dynamics (fig. S5) in 1852 MSNs in the nucleus accumbens core (NAcore, the ventral extension of the caudate-putamen) *in vivo* (fig. S6A and B) during locomotor responses to cocaine in cross-sensitized (892 cells) and mono-sensitized (960 cells) mice. Unsupervised k-means clustering (fig. S6C and E) revealed the underlying response structure of D1- and D2-MSNs in mono-sensitized mice. Clusters with similar patterns over the recording session were combined and classified as *unchanged,* showing stable activity before and after i.p. cocaine and *inactivated* cells showing depression of Ca^2+^ events in response to cocaine (Fig. 2A). In addition to *unchanged* and *inactivated* clusters, in cross-sensitized mice we also found a unique subpopulation of both D1- and D2-MSNs that were *activated* by an acute i.p. cocaine injection (fig. S6D and F, Fig. 2B).

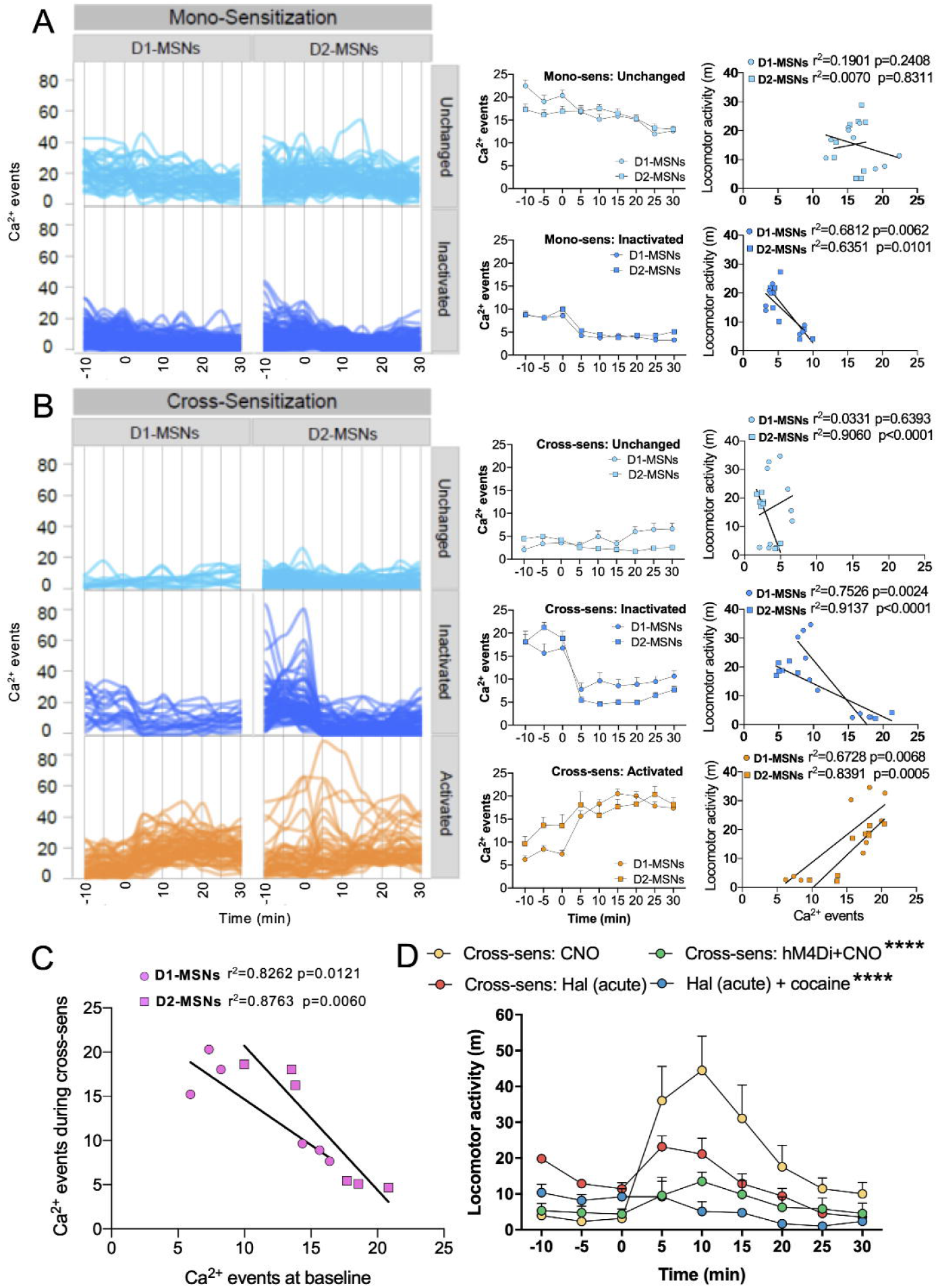

While baseline MSN activity was grossly unchanged in mono-sensitized animals across treatments, the baseline activity of D2-MSNs was enhanced relative to control conditions during ongoing chronic haloperidol treatment (fig. S7). This effect endured after treatment discontinuation. Furthermore, baseline MSN activity after haloperidol discontinuation served as a reliable predictor of cellular responses to cocaine during cross-sensitization (Fig. 2C), in that higher baseline activity predicted depression of cellular response to cocaine whereas moderate basal D2-MSN activity predicted cocaine-induced hyperactivity (as in Fig. 2B, middle panel). When cell clusters were examined separately regarding their relationship to locomotor output, we found that unchanged and decreased MSNs in mono-sensitized (Fig. 2A, right panel) as well as cross-sensitized mice either did not correlate or correlated negatively with locomotion (Fig. 2B, right panel). Instead, both D1- and D2-MSNs that were activated by cocaine in cross-sensitized animals correlated positively with locomotion, suggesting an important role for hyperactive MSN subpopulations in mediating behavioral supersensitivity (Fig. 2B, right panel). Importantly, locomotor activation by psychostimulants is thought to stem from the combined stimulation of D1-and D2-MSNs (*37*). Here instead we found that both D1- and D2-MSNs show similar patterns of responding to cocaine in mono- and cross-sensitized animals and that subpopulations of D2-MSNs were uniquely activated by cocaine only in cross-sensitized animals following haloperidol discontinuation (Fig. 2B, *activated*).

To determine the role of hyperactive D2-MSNs in locomotion during behavioral supersensitivity, we next assessed if silencing D2-MSNs could suppress locomotor cross-sensitization. We virally delivered a Gi-coupled designer receptor activated by the designer drug (DREADD) clozapine-N-oxide (CNO) (*38*) to D2-MSNs in the NAcore of haloperidol pretreated mice. Mice that received CNO intracranially to inhibit D2-MSN activity did not exhibit locomotor cross-sensitization after cocaine injection compared with mice not expressing the Gi-DREADD (Fig. 2D). Locomotion in mice undergoing DREADD inhibition in D2-MSNs was comparable to locomotion in untreated mice that received acute haloperidol, which suppressed cocaine-induced locomotion. We also found that acute haloperidol in animals that had undergone treatment discontinuation was not sufficient to block cocaine-induced hyperlocomotion (Fig. 2D), consistent with loss of antipsychotic efficacy (*5, 7*) and in keeping with emergence of refractory motor symptoms during behavioral supersensitivity. Altogether these findings show that hyperactive D2-MSNs mediate behavioral supersensitivity. Furthermore, since acute haloperidol only partly suppressed hyperlocomotion during cross-sensitization we confirm that D2r blockade is functionally unrelated to the contribution of D2-MSN activity on motor output (*39*).

### Enhanced excitatory transmission on D2-MSNs underlies behavioral supersensitivity

NAcore MSN excitability is regulated by cortical, thalamic and limbic excitatory inputs, and D2-MSNs likely receive specific inputs from the cortex (*40, 41*). Additionally, NAcore MSN excitability is modulated by astrocytes via their expression of the glutamate transporter GLT-1 that regulates glutamate uptake following synaptic release (*42*) and their synaptic proximity, a dynamic measure that impacts autoinhibitory control of transmitter release (*43, 44*). Moreover, striatal glutamatergic transmission is altered by antipsychotic drugs (*15*) and may involve modification of recycling and readily-releasable synaptic vesicle mechanisms (*15, 45*). To determine whether density of presynaptic vesicles was impacted by haloperidol during and after treatment discontinuation, we immunolabeled tissue from haloperidol-treated mice for the presynaptic marker Synapsin I. We found increased density of Synapsin I-positive puncta in the NAcore during chronic haloperidol treatment and discontinuation (Fig. 3D), indicating presynaptic changes capable of elevating transmitter release capacity. Next, we labeled NAcore astroglia using a virally-delivered membrane-bound fluorophore (mCherry) and measured changes in synaptic proximity of NAcore astroglia by quantifying confocal co-registration of mCherry with immunoreactive Synapsin I. Synaptic co-registration of the astroglial membrane was reduced after chronic haloperidol treatment and during cross-sensitization (Fig. 3E). These data illustrate synaptic retraction of astrocyte processes during behavioral supersensitivity, an adaptation that would be expected to reduce autoinhibitory control of transmitter release. We next immunolabeled GLT-1 on NAcore astroglia and found increased GLT-1 expression during haloperidol treatment that returned to baseline levels during abstinence (Fig. 3F), despite persistent increases in Synapsin I labeling (Fig. 3D). Together, these adaptations along with reduced synaptic proximity of astroglial processes that harbor GLT-1 (Fig. 3E) would be expected to render synapses vulnerable to overexcitation during cross-sensitization.

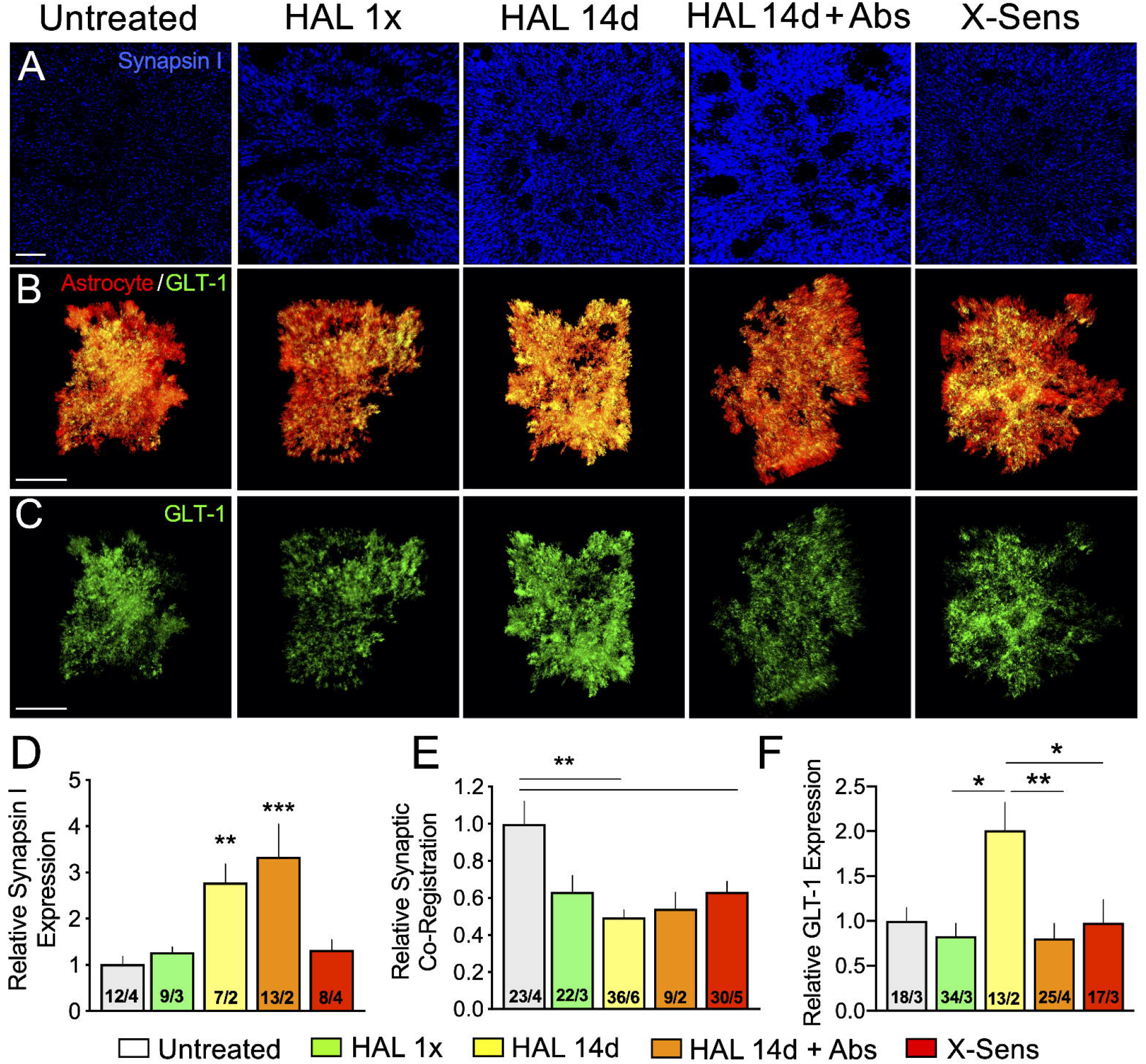

### Neurobiology of behavioral supersensitivity

We show for the first time a crucial role for NAcore D2-MSN activation in driving locomotor activity during behavioral supersensitivity. Beyond the relevance to treatment for schizophrenia, these data invite us to rethink the canonical role of striatal D2-MSN physiology underlying motor outputs. It is generally thought that stimulation of D1 and D2 receptors with either dopamine or with direct and indirect agonists produce stimulation and inhibition of MSNs via the activation of intracellular G_s/o_ or G_i/o_ proteins, respectively. On the contrary we show not only that D1-MSNs are not activated by cocaine during mono-sensitization (Fig. 2A), but also that silencing D2-MSNs inhibits locomotor activity in mice discontinuing antipsychotic treatment during cross-sensitization, revealing a subpopulation of D2-MSNs that can be triggered to drive hyperlocomotion. It is important to note that when MSN activity during cross-sensitization was averaged, D1- and D2-MSNs were increased and decreased in response to cocaine, respectively, in line with canonical expectation of the impact of cocaine on MSN activity (fig. S8). This “canonical” outcome was not observed in the mono-sensitization group. Moreover, an important and opposing role for D2-MSN activation on locomotion was uncovered by teasing apart activity in cell clusters. D2-MSN inhibition in this case suppressed locomotion, consistent with a functional role of hyperactive D2-MSNs in driving locomotion during behavioral supersensitivity.

### Relevance for antipsychotic treatment in human patients

While side effects of antipsychotic treatment are often attributed to changes in the D2r, our study shows that antipsychotic discontinuation leads to the emergence of behavioral supersensitivity, refractory response to antipsychotic treatment and addiction vulnerability independent from D2r changes. Most importantly we show evidence for long-term adaptations driving hyperactivated MSNs during psychotic-like behaviors after antipsychotic discontinuation. These studies prove that antipsychotic discontinuation itself, in the absence of schizophrenia, contributes to endophenotypes of substance use disorder. Deficits in behavioral extinction and increased reinstated seeking would be expected to translate to increased relapse rates and shorter abstinence periods in humans undergoing parallel pharmacological treatments. Since D2-MSNs have been shown to mediate both aversion and reward (*46*), increased lever pressing during extinction and reinstatement may derive from the motivation of subjects to either reduce aversion during antipsychotic discontinuation or to achieve reward.

## ACKNOWLEDGEMENTS

We thank Mr. Eric Dereschewitz as well as Drs. Qing Liu, Alexia Thomas, Rusty Nall, Jasper Heinsbroek and Pete Vento for technical assistance. We also thank Dr. Peter Kalivas for MATLAB codes and experimental animals and both Dr. Kalivas and Dr. Tom Jhou for lab space. Funding: D.A. and the entire study were supported by Deutsche Forschungsgemeinschaft (AM 488 /1-1/-2) and the Brain & Behavior Research Foundation. J.P.C. was supported by the NIH (DA037327 award to Dr. Tom Jhou). Author contributions: D.A. designed the experiments. D.A., A.K., J.P.C., C.P., A.A., R.M.C. performed the behavioral and neurobiological studies with input from S.R. L.B. did the meta-analysis. D.A. and A.K. analyzed the data and drafted the manuscript and all authors edited the final version of the manuscript. Competing interests: The authors declare no competing interests. Data and materials availability: All experimental data are available in the main text or within the supplement.

---

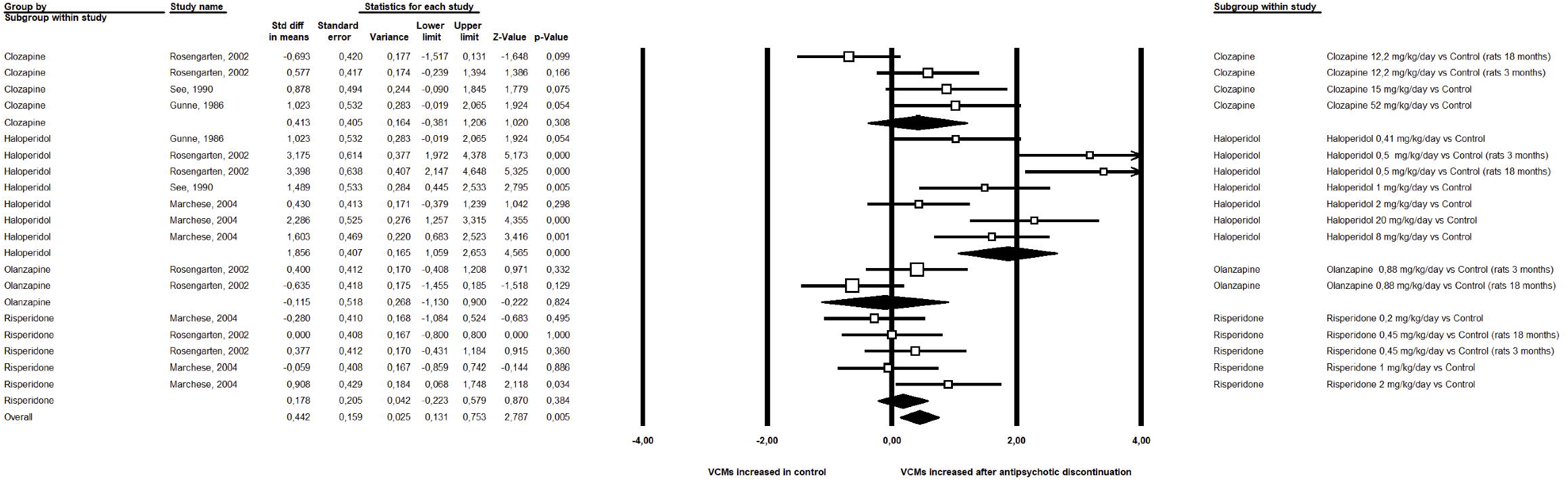

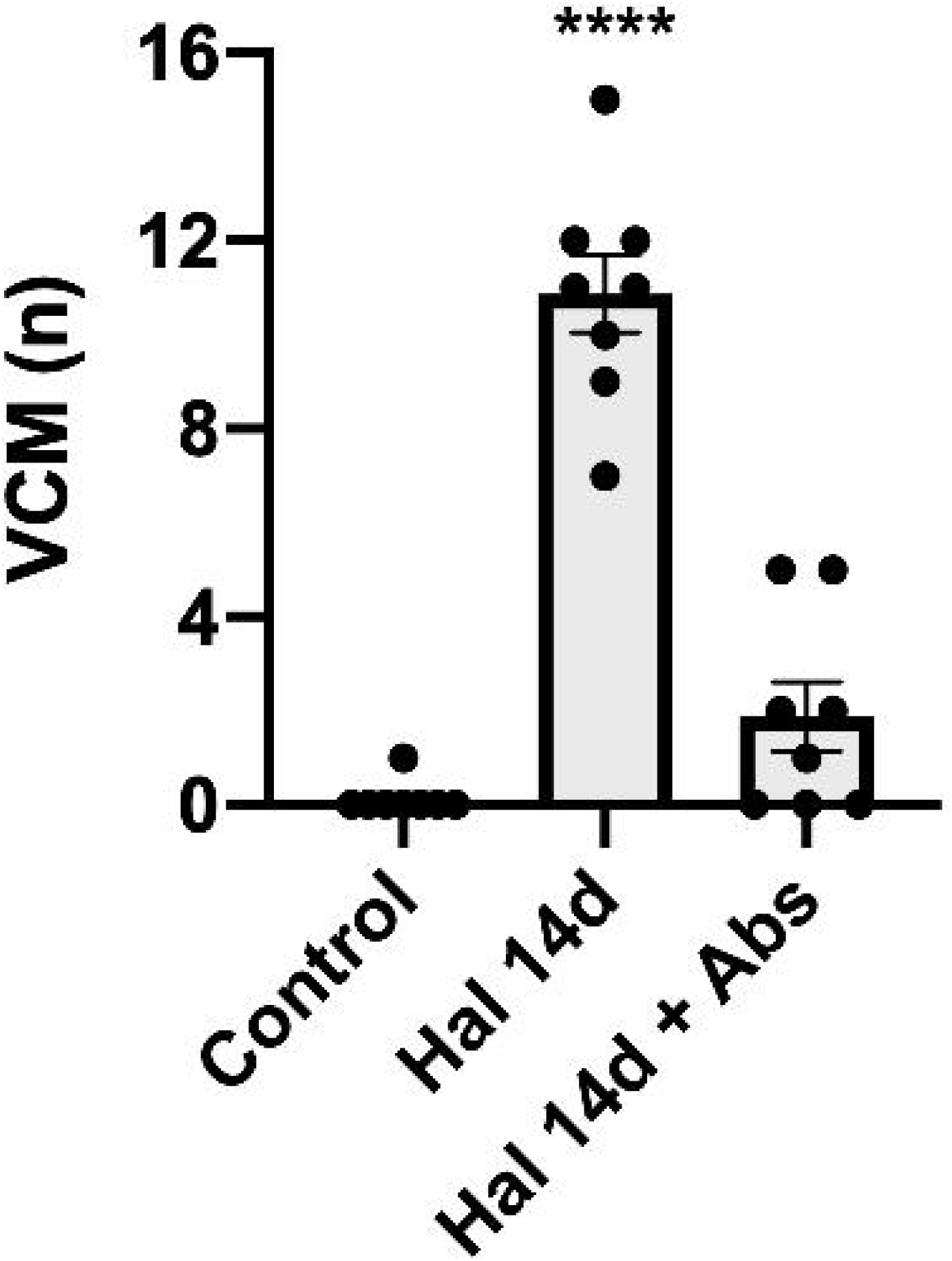

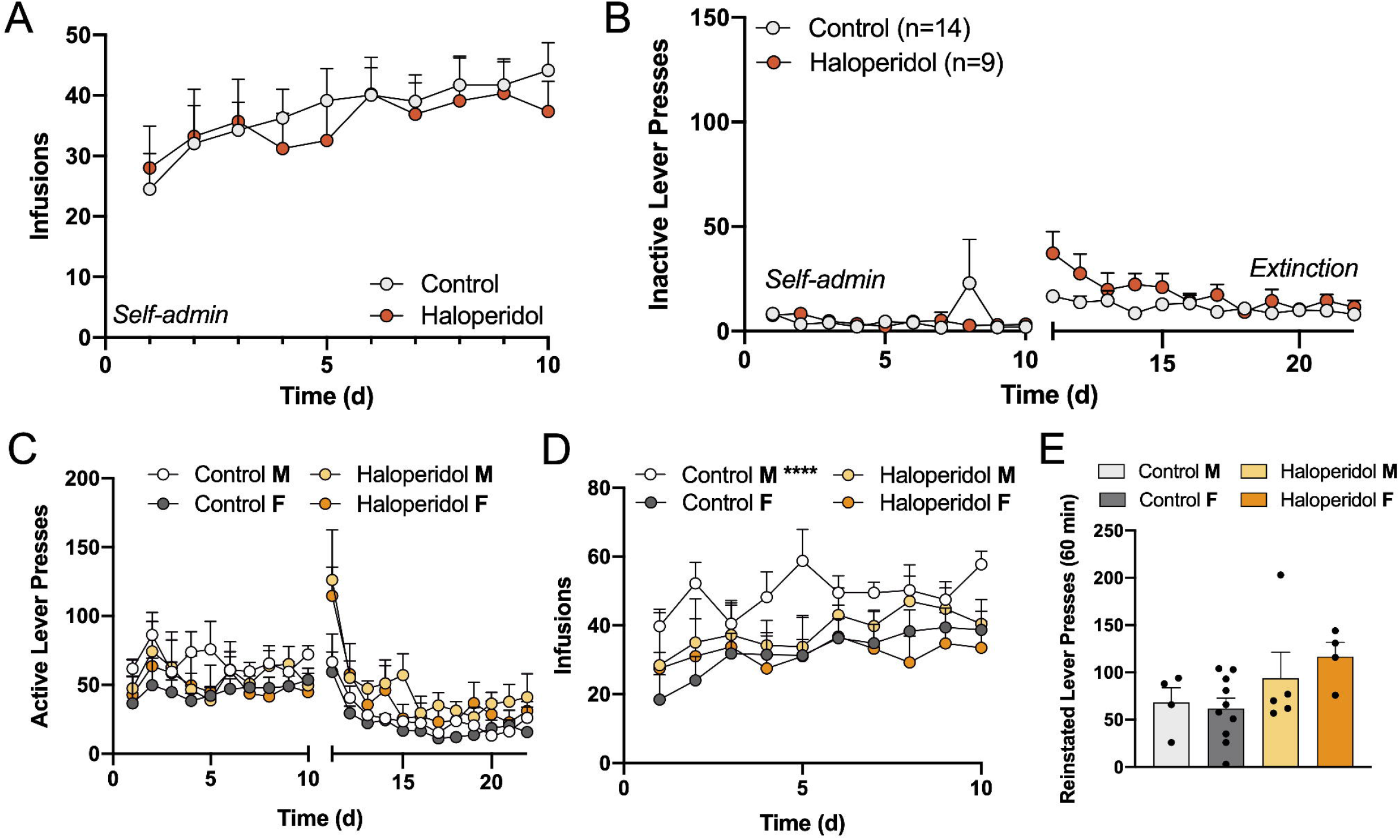

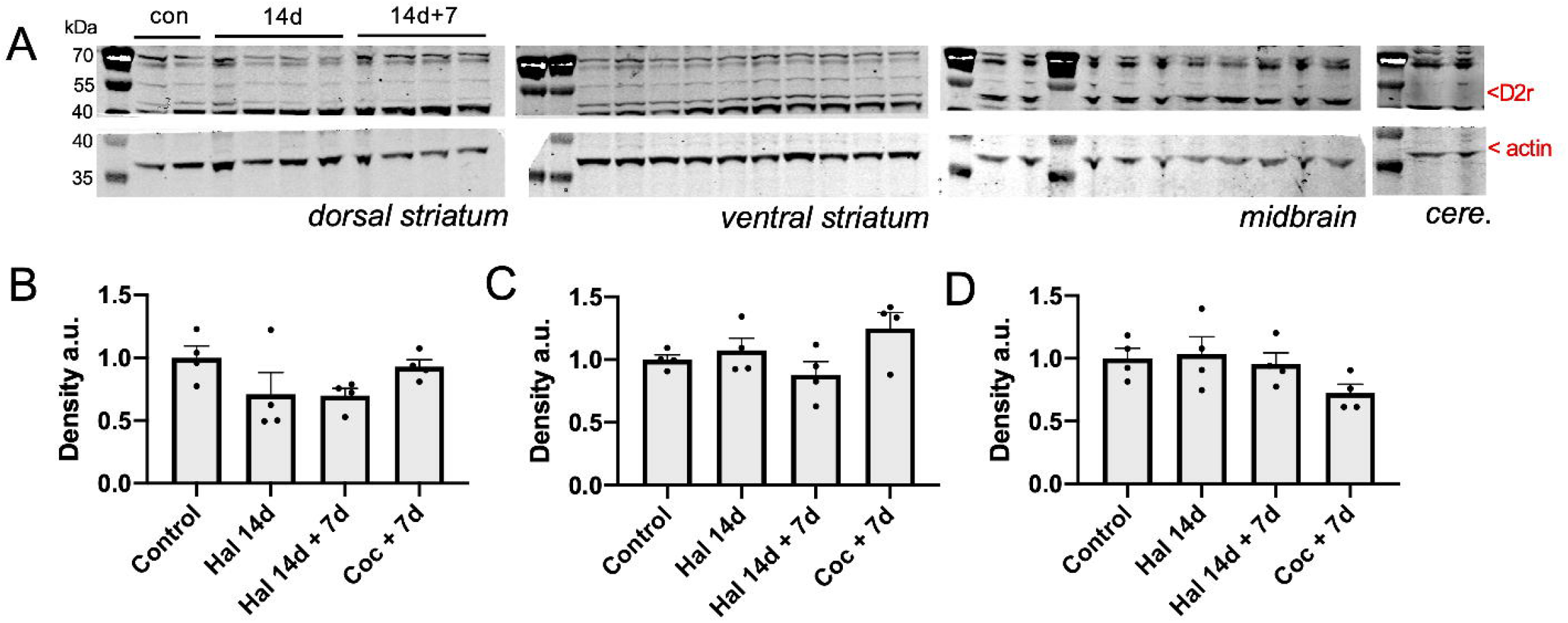

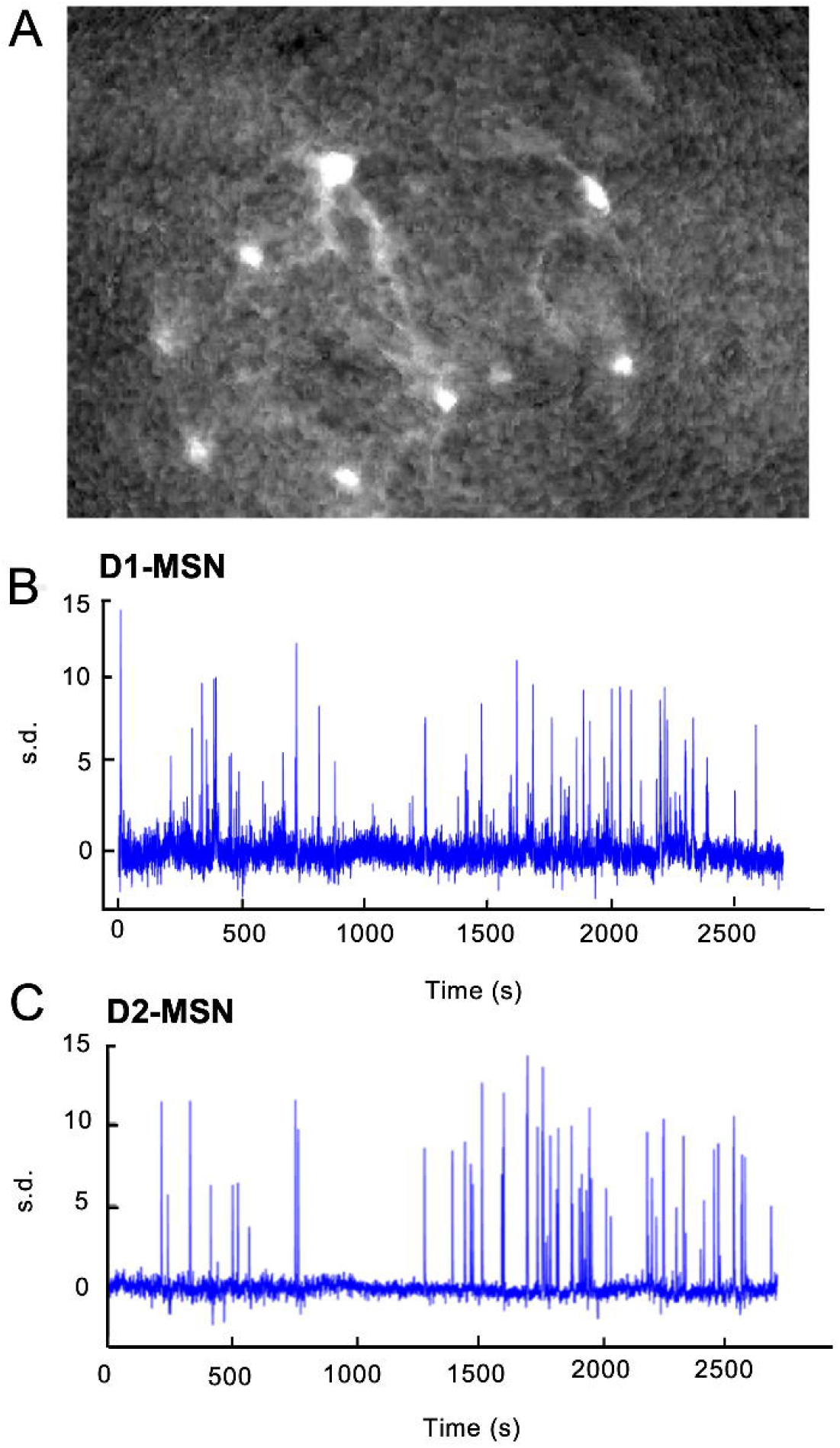

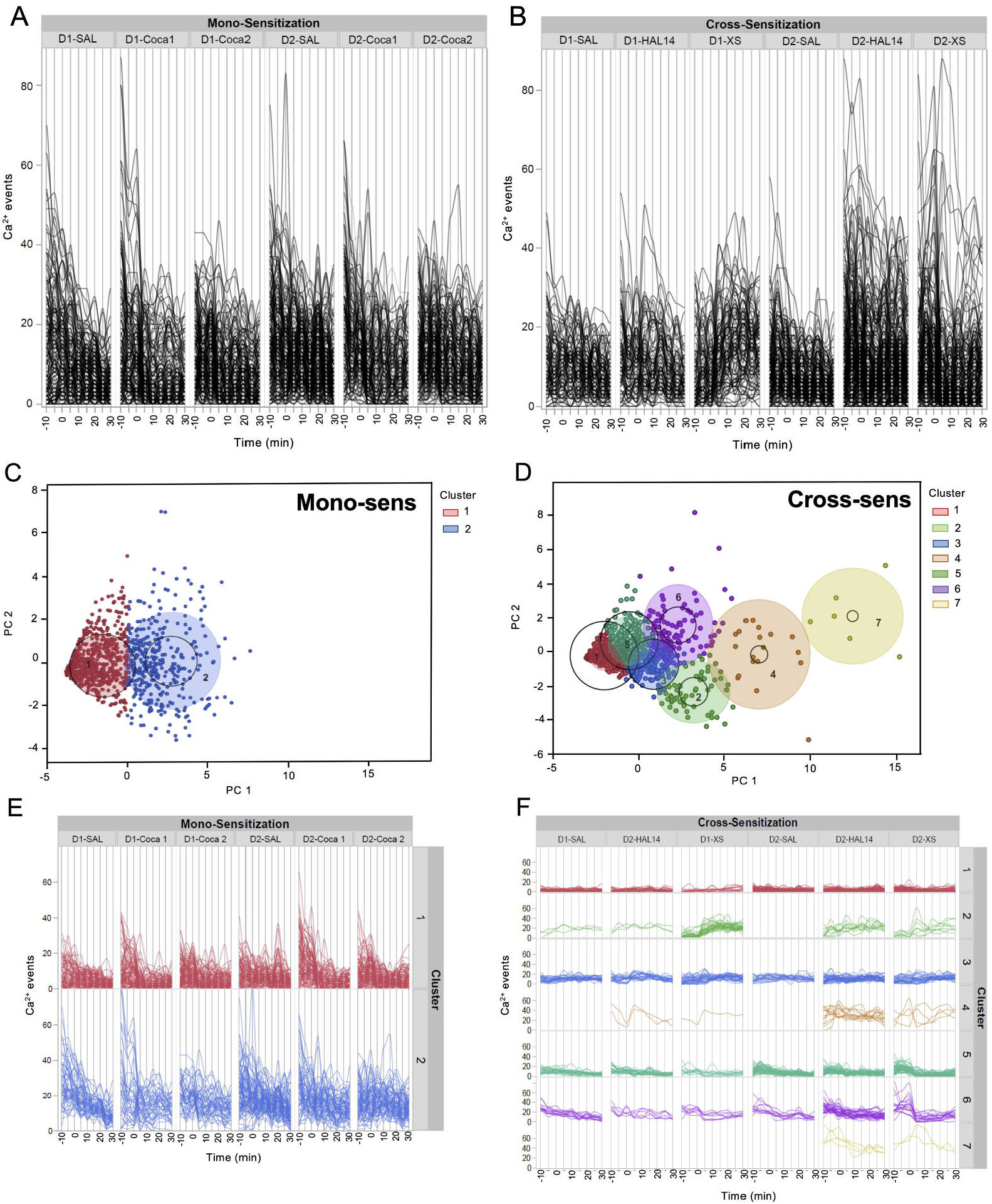

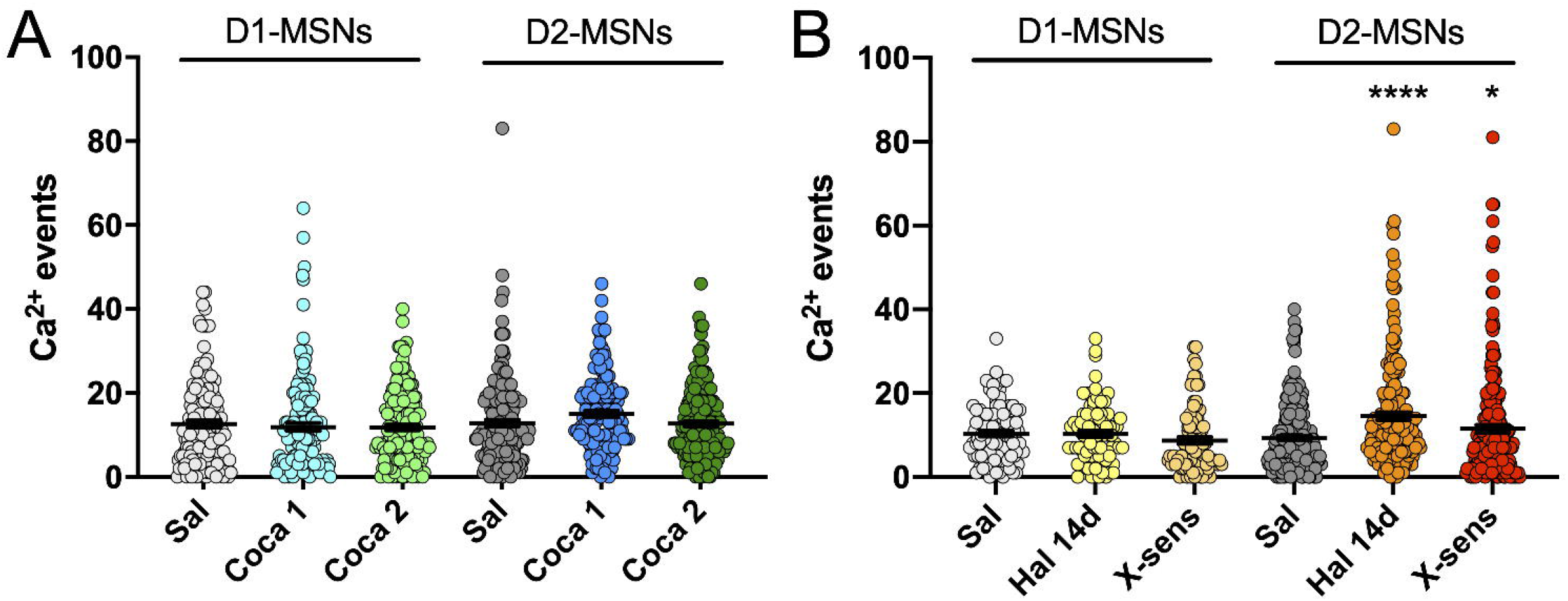

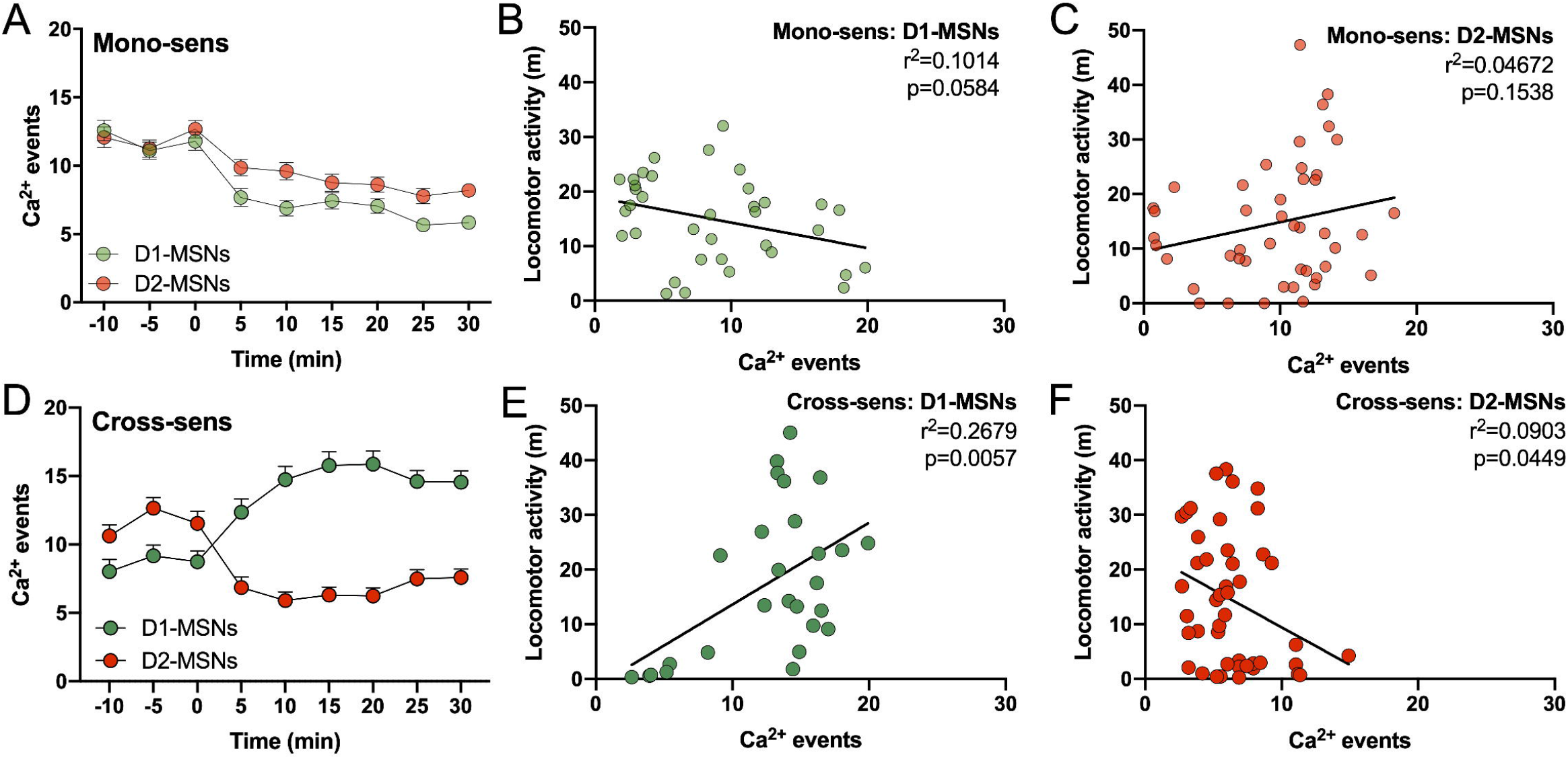

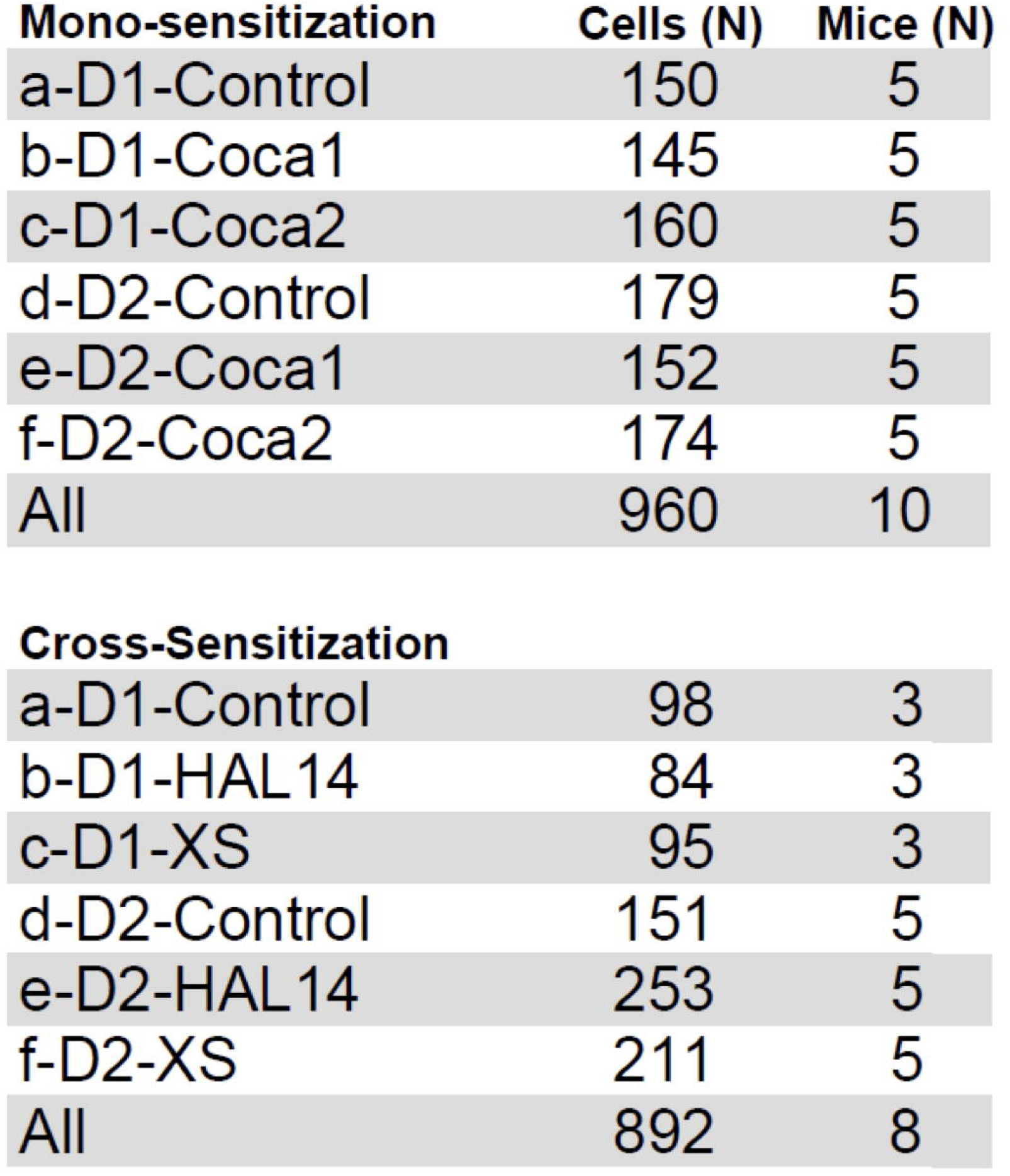

## Notes

### Competing Interest Statement

The authors have declared no competing interest.

