## Supplement for "Accumbens D2-MSN hyperactivity drives behavioral supersensitivity"

**Supplementary information**

To find studies on animal experimentation in PubMed we used the search filter described by Hooijmans et al. (2010) (*47*), the antipsychotics classification described in the WHO ATC classification system (*52*), and terms for withdrawal in the search terms:

(("animal experimentation"[MeSH Terms] OR "models, animal"[MeSH Terms] OR "invertebrates"[MeSH Terms] OR "Animals"[Mesh:noexp] OR "animal population groups"[MeSH Terms] OR "chordata"[MeSH Terms:noexp] OR "chordata, nonvertebrate"[MeSH Terms] OR "vertebrates"[MeSH Terms:noexp] OR "amphibians"[MeSH Terms] OR "birds"[MeSH Terms] OR "fishes"[MeSH Terms] OR "reptiles"[MeSH Terms] OR "mammals"[MeSH Terms:noexp] OR "primates"[MeSH Terms:noexp] OR "artiodactyla"[MeSH Terms] OR "carnivora"[MeSH Terms] OR "cetacea"[MeSH Terms] OR "chiroptera"[MeSH Terms] OR "elephants"[MeSH Terms] OR "hyraxes"[MeSH Terms] OR "insectivora"[MeSH Terms] OR "lagomorpha"[MeSH Terms] OR "marsupialia"[MeSH Terms] OR "monotremata"[MeSH Terms] OR "perissodactyla"[MeSH Terms] OR "rodentia"[MeSH Terms] OR "scandentia"[MeSH Terms] OR "sirenia"[MeSH Terms] OR "xenarthra"[MeSH Terms] OR "haplorhini"[MeSH Terms:noexp] OR "strepsirhini"[MeSH Terms] OR "platyrrhini"[MeSH Terms] OR "tarsii"[MeSH Terms] OR "catarrhini"[MeSH Terms:noexp] OR "cercopithecidae"[MeSH Terms] OR "hylobatidae"[MeSH Terms] OR "hominidae"[MeSH Terms:noexp] OR "gorilla gorilla"[MeSH Terms] OR "pan paniscus"[MeSH Terms] OR "pan troglodytes"[MeSH Terms] OR "pongo pygmaeus"[MeSH Terms])

OR

((animals[tiab] OR animal[tiab] OR mice[Tiab] OR mus[Tiab] OR mouse[Tiab] OR murine[Tiab] OR woodmouse[tiab] OR rats[Tiab] OR rat[Tiab] OR murinae[Tiab] OR muridae[Tiab] OR cottonrat[tiab] OR cottonrats[tiab] OR hamster[tiab] OR hamsters[tiab] OR cricetinae[tiab] OR rodentia[Tiab] OR rodent[Tiab] OR rodents[Tiab] OR pigs[Tiab] OR pig[Tiab] OR swine[tiab] OR swines[tiab] OR piglets[tiab] OR piglet[tiab] OR boar[tiab] OR boars[tiab] OR "sus scrofa"[tiab] OR ferrets[tiab] OR ferret[tiab] OR polecat[tiab] OR polecats[tiab] OR "mustela putorius"[tiab] OR "guinea pigs"[Tiab] OR "guinea pig"[Tiab] OR cavia[Tiab] OR callithrix[Tiab] OR marmoset[Tiab] OR marmosets[Tiab] OR cebuella[Tiab] OR hapale[Tiab] OR octodon[Tiab] OR chinchilla[Tiab] OR chinchillas[Tiab] OR gerbillinae[Tiab] OR gerbil[Tiab] OR gerbils[Tiab] OR jird[Tiab] OR jirds[Tiab] OR merione[Tiab] OR meriones[Tiab] OR rabbits[Tiab] OR rabbit[Tiab] OR hares[Tiab] OR hare[Tiab] OR diptera[Tiab] OR flies[Tiab] OR fly[Tiab] OR dipteral[Tiab] OR drosphila[Tiab] OR drosophilidae[Tiab] OR cats[Tiab] OR cat[Tiab] OR carus[Tiab] OR felis[Tiab] OR nematoda[Tiab] OR nematode[Tiab] OR nematoda[Tiab] OR nematode[Tiab] OR nematodes[Tiab] OR sipunculida[Tiab] OR dogs[Tiab] OR dog[Tiab] OR canine[Tiab] OR canines[Tiab] OR canis[Tiab] OR sheep[Tiab] OR sheeps[Tiab] OR mouflon[Tiab] OR mouflons[Tiab] OR ovis[Tiab] OR goats[Tiab] OR goat[Tiab] OR capra[Tiab] OR capras[Tiab] OR rupicapra[Tiab] OR chamois[Tiab] OR haplorhini[Tiab] OR monkey[Tiab] OR monkeys[Tiab] OR anthropoidea[Tiab] OR anthropoids[Tiab] OR saguinus[Tiab] OR tamarin[Tiab] OR tamarins[Tiab] OR leontopithecus[Tiab] OR hominidae[Tiab] OR ape[Tiab] OR apes[Tiab] OR pan[Tiab] OR paniscus[Tiab] OR "pan paniscus"[Tiab] OR bonobo[Tiab] OR bonobos[Tiab] OR troglodytes[Tiab] OR "pan troglodytes"[Tiab] OR gibbon[Tiab] OR gibbons[Tiab] OR siamang[Tiab] OR siamangs[Tiab] OR nomascus[Tiab] OR symphalangus[Tiab] OR chimpanzee[Tiab] OR chimpanzees[Tiab] OR prosimians[Tiab] OR "bush baby"[Tiab] OR prosimian[Tiab] OR bush babies[Tiab] OR galagos[Tiab] OR galago[Tiab] OR pongidae[Tiab] OR gorilla[Tiab] OR gorillas[Tiab] OR pongo[Tiab] OR pygmaeus[Tiab] OR "pongo pygmaeus"[Tiab] OR orangutans[Tiab] OR pygmaeus[Tiab] OR lemur[Tiab] OR lemurs[Tiab] OR lemuridae[Tiab] OR horse[Tiab] OR horses[Tiab] OR pongo[Tiab] OR equus[Tiab] OR cow[Tiab] OR calf[Tiab] OR bull[Tiab] OR chicken[Tiab] OR chickens[Tiab] OR gallus[Tiab] OR quail[Tiab] OR bird[Tiab] OR birds[Tiab] OR quails[Tiab] OR poultry[Tiab] OR poultries[Tiab] OR fowl[Tiab] OR fowls[Tiab] OR reptile[Tiab] OR reptilia[Tiab] OR reptiles[Tiab] OR snakes[Tiab] OR snake[Tiab] OR lizard[Tiab] OR lizards[Tiab] OR alligator[Tiab] OR alligators[Tiab] OR crocodile[Tiab] OR crocodiles[Tiab] OR turtle[Tiab] OR turtles[Tiab] OR amphibian[Tiab] OR amphibians[Tiab] OR amphibia[Tiab] OR frog[Tiab] OR frogs[Tiab] OR bombina[Tiab] OR salientia[Tiab] OR toad[Tiab] OR toads[Tiab] OR "epidalea calamita"[Tiab] OR salamander[Tiab] OR salamanders[Tiab] OR eel[Tiab] OR eels[Tiab] OR fish[Tiab] OR fishes[Tiab] OR pisces[Tiab] OR catfish[Tiab] OR catfishes[Tiab] OR siluriformes[Tiab] OR arius[Tiab] OR heteropneustes[Tiab] OR sheatfish[Tiab] OR perch[Tiab] OR perches[Tiab] OR percidae[Tiab] OR perca[Tiab] OR trout[Tiab] OR trouts[Tiab] OR char[Tiab] OR chars[Tiab] OR salvelinus[Tiab] OR "fathead minnow"[Tiab] OR minnow[Tiab] OR cyprinidae[Tiab] OR carps[Tiab] OR carp[Tiab] OR zebrafish[Tiab] OR zebrafishes[Tiab] OR goldfish[Tiab] OR goldfishes[Tiab] OR guppy[Tiab] OR guppies[Tiab] OR chub[Tiab] OR chubs[Tiab] OR tinca[Tiab] OR barbels[Tiab] OR barbus[Tiab] OR pimephales[Tiab] OR promelas[Tiab] OR "poecilia reticulata"[Tiab] OR mullet[Tiab] OR mullets[Tiab] OR seahorse[Tiab] OR seahorses[Tiab] OR mugil curema[Tiab] OR atlantic cod[Tiab] OR shark[Tiab] OR sharks[Tiab] OR catshark[Tiab] OR anguilla[Tiab] OR salmonid[Tiab] OR salmonids[Tiab] OR whitefish[Tiab] OR whitefishes[Tiab] OR salmon[Tiab] OR salmons[Tiab] OR sole[Tiab] OR solea[Tiab] OR "sea lamprey"[Tiab] OR lamprey[Tiab] OR lampreys[Tiab] OR pumpkinseed[Tiab] OR sunfish[Tiab] OR sunfishes[Tiab] OR tilapia[Tiab] OR tilapias[Tiab] OR turbot[Tiab] OR turbots[Tiab] OR flatfish[Tiab] OR flatfishes[Tiab] OR sciuridae[Tiab] OR squirrel[Tiab] OR squirrels[Tiab] OR chipmunk[Tiab] OR chipmunks[Tiab] OR suslik[Tiab] OR susliks[Tiab] OR vole[Tiab] OR voles[Tiab] OR lemming[Tiab] OR lemmings[Tiab] OR muskrat[Tiab] OR muskrats[Tiab] OR lemmus[Tiab] OR otter[Tiab] OR otters[Tiab] OR marten[Tiab] OR martens[Tiab] OR martes[Tiab] OR weasel[Tiab] OR badger[Tiab] OR badgers[Tiab] OR ermine[Tiab] OR mink[Tiab] OR minks[Tiab] OR sable[Tiab] OR sables[Tiab] OR gulo[Tiab] OR gulos[Tiab] OR wolverine[Tiab] OR wolverines[Tiab] OR minks[Tiab] OR mustela[Tiab] OR llama[Tiab] OR llamas[Tiab] OR alpaca[Tiab] OR alpacas[Tiab] OR camelid[Tiab] OR camelids[Tiab] OR guanaco[Tiab] OR guanacos[Tiab] OR chiroptera[Tiab] OR chiropteras[Tiab] OR bat[Tiab] OR bats[Tiab] OR fox[Tiab] OR foxes[Tiab] OR iguana[Tiab] OR iguanas[Tiab] OR xenopus laevis[Tiab] OR parakeet[Tiab] OR parakeets[Tiab] OR parrot[Tiab] OR parrots[Tiab] OR donkey[Tiab] OR donkeys[Tiab] OR mule[Tiab] OR mules[Tiab] OR zebra[Tiab] OR zebras[Tiab] OR shrew[Tiab] OR shrews[Tiab] OR bison[Tiab] OR bisons[Tiab] OR buffalo[Tiab] OR buffaloes[Tiab] OR deer[Tiab] OR deers[Tiab] OR bear[Tiab] OR bears[Tiab] OR panda[Tiab] OR pandas[Tiab] OR "wild hog"[Tiab] OR "wild boar"[Tiab] OR fitchew[Tiab] OR fitch[Tiab] OR beaver[Tiab] OR beavers[Tiab] OR jerboa[Tiab] OR jerboas[Tiab] OR capybara[Tiab] OR capybaras[Tiab]) NOT medline[subset])

AND

((Acepromazine[tiab] OR Acetophenazine[tiab] OR Benperidol[tiab] OR Bromperidol[tiab] OR Butaperazine[tiab] OR Carfenazine[tiab] OR Chlorproethazine[tiab] OR Chlorpromazine[tiab] OR Chlorprothixene[tiab] OR Clopenthixol[tiab] OR Cyamemazine[tiab] OR Dixyrazine[tiab] OR Droperidol[tiab] OR Fluanisone[tiab] OR Flupentixol[tiab] OR Fluphenazine[tiab] OR Fluspirilene[tiab] OR Haloperidol[tiab] OR Levomepromazine[tiab] OR Lenperone[tiab] OR Loxapine[tiab] OR Mesoridazine[tiab] OR Metitepine[tiab] OR Molindone[tiab] OR Moperone[tiab] OR Oxypertine[tiab] OR Oxyprotepine[tiab] OR Penfluridol[tiab] OR Perazine[tiab] OR Periciazine[tiab] OR Perphenazine[tiab] OR Pimozide[tiab] OR Pipamperone[tiab] OR Piperacetazine[tiab] OR Pipotiazine[tiab] OR Prochlorperazine[tiab] OR Promazine[tiab] OR Prothipendyl[tiab] OR Spiperone[tiab] OR Sulforidazine[tiab] OR Thiopropazate[tiab] OR Thioproperazine[tiab] OR Thioridazine[tiab] OR Thiothixene[tiab] OR Timiperone[tiab] OR Trifluoperazine[tiab] OR Trifluperidol[tiab] OR Triflupromazine[tiab] OR Zuclopenthixol[tiab] OR Amoxapine[tiab] OR Amisulpride[tiab] OR Aripiprazole[tiab] OR Asenapine[tiab] OR Blonanserin[tiab] OR Brexpiprazole[tiab] OR Cariprazine[tiab] OR Carpipramine[tiab] OR Clocapramine[tiab] OR Clorotepine[tiab] OR Clotiapine[tiab] OR Clozapine[tiab] OR Iloperidone[tiab] OR Levosulpiride[tiab] OR Lurasidone[tiab] OR Melperone[tiab] OR Mosapramine[tiab] OR Nemonapride[tiab] OR Olanzapine[tiab] OR Paliperidone[tiab] OR Perospirone[tiab] OR Quetiapine[tiab] OR Remoxipride[tiab] OR Reserpine[tiab] OR Risperidone[tiab] OR Sertindole[tiab] OR Sulpiride[tiab] OR Sultopride[tiab] OR Tiapride[tiab] OR Veralipride[tiab] OR Ziprasidone[tiab] OR Zotepine[tiab] OR "antipsychotic agents"[MeSH Terms] antipsychotic*[tiab] OR neuroleptic*[tiab])

AND

("substance withdrawal syndrome"[MeSH Terms] OR discontinu*[tiab] OR withdraw*[tiab] OR ceas*[tiab] OR withhold*[tiab] OR stop*[tiab] OR end[tiab] OR ended[tiab] OR ending[tiab])))

**PRISMA flowchart**

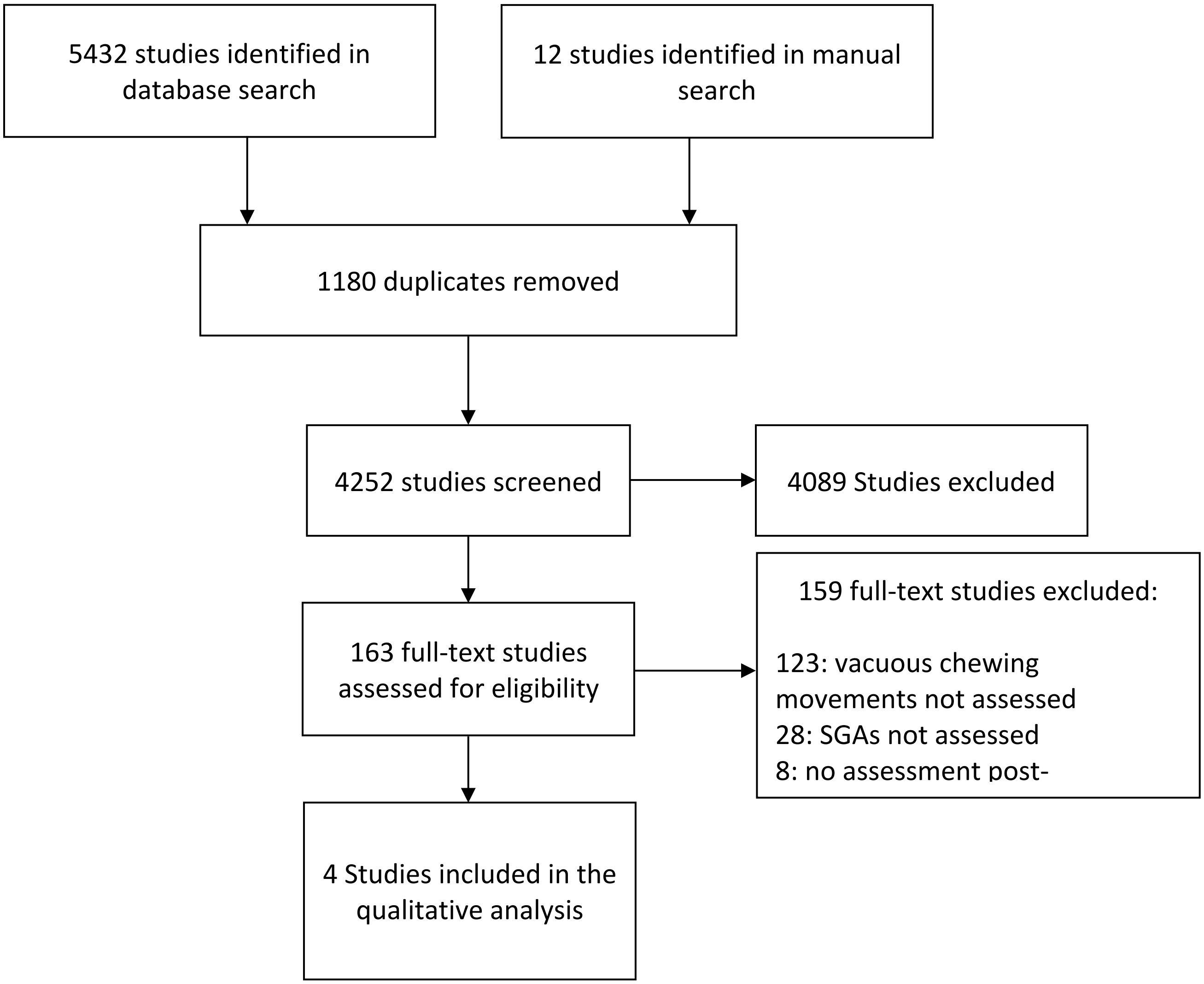

**METHODS**

**Meta-analysis.**

We searched PubMed, EMBASE, and Web of Science for studies until April 1, 2020. No restriction was implemented for the beginning of the searched time period. In addition, we manually searched references from included and relevant reviews. Search terms for (1) animal studies, (2) antipsychotics, and (3) withdrawal were used for filtering studies (Supplementary information). We included antipsychotics classified in The Anatomical Therapeutic Chemical (ATC) classification system by the WHO Collaborating Centre for Drug Statistics Methodology[1] in the search terms. For PubMed and Web of Science we included the search filter for finding studies on animal experimentation by Hooijmans et al. (2010) (*47*) and for EMBASE we included the updated version of the EMBASE search filter for animal studies by de Vries et al. (2014) (*48*). Search terms by Hooijmans et al. (2010) and de Vries et al. (2014) were recently implemented in a publication by McCann et al. (2020) (*49*) for searching animal models of ischaemic stroke. We screened title, abstracts, and key words and implemented no language restrictions. The literature screening of title, abstract, and key words was carried out by one researcher and testing of eligibility criteria in full-text were carried out independently by two researchers. Discrepancies were resolved by consensus with an additional researcher from the review team. Authors of the original articles were contacted if information was missing in the original articles, provided contact information could be retrieved. The software Endnote, version X8.2 (Clarivate Analytics) was used for the literature search. Data was included if the behavioural assessments of VCMs in animal models were employed after discontinuation of haloperidol and any SGA (*50*). Studies in humans, only single applications of antipsychotics (i.e. not repeated applications of antipsychotics), and studies only on FGAs were excluded. A predefined Excel spreadsheet (Microsoft Excel for Mac, version 16.12, Microsoft Corporation) was used for data collection and extraction from selected studies and data was synthesized quantitatively. We calculated standardized mean differences (SMD) and 95% confidence intervals (CIs) from outcome measures of the primary studies. If respective measures of dispersion were not available, we calculated CIs from $p$ values as recommended in the Cochrane Handbook (*51*) . Stratified according to the type of antipsychotic, effect sizes for comparisons between animals treated with antipsychotics and animals treated with placebo (i.e. control) were summarized using a forest plot. We calculated summary estimates using random-effects models, as the studies differed in several methodological aspects (*51*). Analyses were conducted according to the Cochrane Collaboration Handbook (*51*) and using Comprehensive Meta-Analysis V3 (Biostat, Engelwood, New Jersey).

**Quantitation of vacuous chewing movements.** Rats were moved into the testing room 1h prior to VCM monitoring and habituated to the testing arena for 5 minutes. VCMs, defined as purposeless mouth opening with or without tongue protrusion as described in (*21*), were quantified for each animal during 2-min of observation and averaged within treatment groups.

**Locomotor sensitization and cross-sensitization.** Male and female 8-13-week-old transgenic mice were group housed (3-5/cage) on a reverse light/dark cycle and had *ad libitum* access to food and water. Mice undergoing cross-sensitization were implanted with subcutaneous osmotic infusion pumps (Alzet) delivering haloperidol at 0.5mg/kg/day for 14 days. Mice were abstinent from haloperidol for an additional 7 days before undergoing locomotor cross-sensitization with cocaine. Pumps were removed 7 days before testing in a subgroup of mice only, since no differences in the behavioral or cellular responses to cocaine in the presence or absence of pumps was observed. Mice undergoing cocaine mono-sensitization received two cocaine injections separated by 7 days incubation as described in (*23*). All mice were habituated in a dimly lit (15 lux) open field apparatus measuring 43 x 43 x 30 cm with mild background noise for 10-min on three consecutive days prior to experimentation. On the last of these 3 days, mice received 0.2 mL physiological saline i.p. to habituate them to physical handling. During experimentation, mice were placed in the open field during 15-min of spontaneous baseline locomotion. Mice were then briefly transferred into the home cage where they received an i.p. injection of cocaine (15mg/kg) before being placed in the locomotor chamber for 30 min of locomotor recording using ANY-maze behavior tracking software (Stoelting).

**Cocaine self-administration.** Male and female Long Evans rats (150-350g) were implanted with subcutaneous pumps delivering haloperidol (0.5 mg/kg/d) for 14 days. On day 14, pumps were removed and animals were fitted with intrajugular catheters under anesthesia using isoflurane. 7 days after recovery from surgery, haloperidol pretreated rats and untreated controls began daily 2h cocaine self-administration sessions, where active lever presses were paired with light and tone cues and cocaine delivery (0.4 mg/kg/infusion). Inactive lever presses had no consequence. After 10 days of self-administration, rats underwent extinction training (2h/d), where cues and cocaine delivery were withheld during active lever pressing. Extinction training continued for 12 days. The following day, rats were returned to the operant chamber and light/tone cue pairings were restored to the active lever, but no cocaine was delivered. Reinstatement was continued for 60m.

***In vivo* $Ca^{2+}$ imaging.** Prior to recording, mice were fitted with a head-mounted camera and habituated to the experimental setting for 10-min on three consecutive days as described previously (*39*). During experimentation, $Ca^{2+}$ recordings were temporally synchronized with locomotor activity in the open field by manually launching simultaneous recordings. Grayscale video was recorded using nVista HD Acquisition Software (v.2, Inscopix) at 15 Hz and with 20% LED power. Following data acquisition, images were downsampled 4x and preprocessed for motion correction and regions of interest corresponding to visually identifiable cell bodies were identified using a built-in automated algorithm (Mosaic v. 1.1.2, Inscopix). $Ca^{2+}$ traces from independent neurons were computed based on their spatial and temporal components identified using principal/independent component analysis (PCA/ICA) and confirmed by an investigator blind to animal identifiers. $Ca^{2+}$ events were included if they were ≥ 6x the median absolute deviation of the input trace data and had a minimum mean decay of 0.1 s. $Ca^{2+}$ events were then transformed into binary values, with amplitudes >0 assigned a value of 1 and amplitude ≤0 assigned a value of 0, to obtain the number of spikes per min. Subsequently the sum of spikes per minute for each cell was used to determine spikes in 5 min bins to obtain a total of 9 5-min bins. The first 3 bins reflect $Ca^{2+}$ events at baseline (15 min) and the remaining 6 bins represent $Ca^{2+}$ events after saline or cocaine injection (30 min). Next, we evaluated binary values of $Ca^{2+}$ events for sphericity under basal conditions and determined distinct cell response patterns after treatment using unsupervised K-means clustering for parametric data, which identifies a set of k seeds (i.e. centroids) and then assigns each data point to the nearest cluster seed. By reiteration the initial seeds were then replaced by the cluster means and the data points were reassigned. The process continued until no further changes occurred in the clusters (SAS statistics). The optimal number of clusters was automatically selected using the fit statistic Cubic Cluster Criterion (SAS statistics). Clusters were categorized empirically as activated, inactivated, or unchanged. Cluster analysis and the relative representations were performed using SAS-JMP statistics.

**DREADD inhibition of D2-MSNs.** Mice were implanted with bilateral cannulae (Plastics One) situated above the NAcore (+1.6 mm AP, ± 1.1 mm ML, -3.0 mm DV) along with osmotic pumps for subcutaneous infusion of chronic haloperidol and intracranial injections of AAV2/FLEX-hM4Di-mCherry (Addgene) or sterile saline at 0.5 μL/hemisphere, 0.05 μL/min, with 5 min diffusion. Virus incubation occurred during 14 days of haloperidol treatment and 7 days of abstinence following treatment (i.e. 3 weeks). Baseline locomotion was recorded for 15-min before animals were briefly returned to the home cage. Clozapine N-oxide (CNO, 1 mM, 0.3 μL, Abcam) was infused intracranially with bilateral microinjectors extending 1.4 mm DV beyond the tip of the guide cannulae 5-min prior to cocaine injection (15 mg/kg i.p.) and behavioral testing. Next mice were placed in the open field apparatus for 30-min of locomotor recording. Additional subjects received acute haloperidol (0.5 mg/kg i.p.) prior to cocaine injection and locomotor recording.

**Confocal microscopy.** 3 weeks before sacrifice, mice received NAcore injections (+1.6 mm AP, +1.1 mm ML, -4.4 mm DV) of a virus used to label the astroglial membrane (AAV5-GFAP-hM3d-mcherry, University of Zurich) under isoflurane anesthesia. Mice were anesthetized using i.p. ketamine and perfused transcardially with 1X phosphate buffer (10 mL) and 4% PFA (20 mL) before brain extraction. Brains were post-fixed for 24h in 4% PFA before slicing at 100 μm using a vibrating blade microtome. Sections containing the NAcore were permeabilized in 1X PBS with 2% Triton X-100 for 1h at room temperature shaking gently. Tissue was then blocked in 1X PBS with 0.2% Triton X-100 (PBST) and 2% normal goat serum (block) for 1h at room temperature before overnight incubation in primary antibodies (1:1000 anti-GLT-1, ab1783, EMD Millipore and 1:1000 anti-Synapsin I, ab64581, Abcam) in block at 4C. After washing in PBST, tissue was incubated overnight in biotinylated anti-guinea pig antibody (1:1000, Vector Labs) at room temperature in PBST then overnight in fluorescently labeled antibodies (Thermo Fisher) at room temperature in PBST after washing. Tissue was then washed in PBST and mounted onto glass slides. NAcore tissue was imaged using an SP5 confocal microscope (Leica) with the following conditions: 1024 x 1024 frame size, 12-bit resolution, 4-frame averaging, 1-μm step size. Z-stacks were iteratively deconvolved (Autoquant) and analyzed for fluorescence intensity using Bitplane Imaris. Total Synapsin I was expression was quantified using the Surface module in full or partial z-stacks and normalized to the frame volume. Rendered astroglia were digitally isolated and their co-registration with Synapsin I was normalized to the astroglial volume and to total Synapsin I signal per equivalent frame size to account for changes in astroglial volume or Synapsin I density. GLT-1 expression that co-registered with the astroglial label was normalized to the astroglial volume. In all cases, data were ultimately normalized to the mean value of the untreated group. Imaging and analyses were conducted blind to animal treatment.

**Western blotting.** Mice were briefly anesthetized before decapitation and brain extraction. Brains were incubated on ice briefly and dorsal striatum, ventral striatum and midbrain tissue extracts were dissected and manually homogenized in ice-cold RIPA buffer with protease and phosphatase inhibitors (Thermo Fisher) for 20 strokes and then sonicated for 60 s. Samples were centrifuged at 10,000 x g for 10 min. Supernatants were isolated and protein content was quantified using a BCA assay (Thermo Fisher). 70 µg of protein was loaded per lane in Criterion Bis-Tris gradient gels (Bio-Rad). Proteins were then transferred onto a PVDF membrane (Bio-Rad) overnight at 4C. Blots were cut at 40 kDa according to the protein ladder (Thermo Fisher) and blocked in Odyssey blocking buffer (LI-COR) for 1h at room temperature before overnight incubation in primary antibody (anti-actin, ab95437, Abcam or anti-D2r, AB5084P, EMD Millipore) in block at 4C. After rinsing, blots were incubated in secondary antibody (LI-COR) at room temperature for 1h and imaged using a LI-COR Odyssey imaging system. The band corresponding to the D2r was identified based on its molecular weight (~50 kDa) as well as its absence in a blot containing tissue extracts from the cerebellum. Images were converted to grayscale and quantified using FIJI.

**Statistics.** Data were analyzed using a Student's t-test or 1- or 2-way ANOVA with repeated measures when possible. Cluster analysis and relative data representations were performed using SAS-JMP statistics. A Pearson's correlation coefficient was calculated to determine the relationship between $Ca^{2+}$ events and locomotion. In all cases, statistical significance was set at $p<0.05$.
